# Towards simple kinetic models of functional dynamics for a kinase subfamily

**DOI:** 10.1101/228528

**Authors:** Mohammad M. Sultan, Gert Kiss, Vijay Pande

## Abstract

Kinases are ubiquitous enzymes involved in the regulation of critical cellular pathways and have been implicated in several cancers. Consequently, the kinetics and thermodynamics of prototypical kinases are of interest and have been the subject of numerous experimental studies. In-silico modeling of the conformational ensembles of these enzymes, on the other hand, is lacking due to inherent computational limitations. Recent algorithmic advances combined with homology modeling and parallel simulations allow us to address this computational sampling bottleneck. Here, we present the results of molecular dynamics (MD) studies for seven Src family kinase (SFK) members Fyn, Lyn, Lck, Hck, Fgr, Yes, and Blk. We present a sequence invariant extension to Markov state models (MSMs), which allows us to quantitatively compare the structural ensembles of the seven kinases. Our findings indicate that in the absence of their regulatory partners, SFK members have similar in-silico dynamics with active state populations ranging from 4-40% and activation timescales in the hundreds of microseconds. Furthermore, we observe several potentially druggable intermediate states, including a pocket next to the ATP binding site that could be potentially targeted via a small molecule inhibitors. These results establish the utility of MSMs for studying protein families.

## Introduction

Protein kinases are important pharmaceutical targets^1,2^ due to their regulatory roles in biochemical pathways in eukaryotic cells^3^. They control a slew of signaling cascades by catalyzing the transfer of γ-phosphate from adenosine triphosphate (ATP) to target substrates^4^. Kinases are thought to account for up to 2% of all encoded genes^5^. Given their physiological role, kinase functionality within cells is tightly monitored.^4^,^6^ Mutations, truncation, and overexpression of kinases have been phenotypically linked to various kinds of cancers^7–9^, viral infections^10^, and other diseases^11^. The kinetics and thermodynamics of several prototypical kinases such as protein kinase A (PKA)^12^, Src^13–16^, Abl^6^,^17^,^18^, EGFR^19^, and Aurora^7^ kinases have been extensively studied by diverse experimental techniques including crystallography^14^,^20^, small angle X-ray scattering(SAXS)^16^, nuclear magnetic resonance(NMR)^2^.

Members of the Src family of tyrosine kinases (SFK) regulate signal transduction of cellular receptors^9^. They are ubiquitously expressed with several established oncogenic truncations and mutations^9^. For example, Fyn kinase is required for proper cell growth^21^. Aberrant Fyn signaling has also been implicated in the pathogenesis of Alzheimer’s disease^11^,^21^ and Dengue virus RNA replication^10^.

While different kinases have been studied via a range of experimental methods, there is a large degree of variance in the kinase sequences studied and the methods employed, which makes it difficult to systemically compare these observables. However, to effectively and precisely target kinases via small molecule inhibitors requires inference of the differences and similarities between their structural, thermodynamics, and kinetic behavior at the ensemble level. Understanding of the SFK’s solution structural ensemble is lacking. For example, human Fgr and Blk have no reported crystal structures, only a single crystal structure has been reported for Fyn^22^, and two for the human Lyn kinase domain. On the other hand, Hck and Src each have close to or more than ten structures available. Such heterogeneity makes it difficult to systemically compare SFK despite their similarities. Members of SFK share a common architecture which includes several regulatory domains^20,23^, connected to the catalytic domain via a linker region. The catalytic domain is functionally active^24^ in the absence of its regulatory partners, and form primary drug targets^14^. Therefore, we chose to focus our efforts on the catalytic domain. The SFK catalytic domain (Figure 1) is primarily composed of a beta-sheet-rich N-lobe and an alpha helical C-lobe. The lobes are connected via the Hinge (Figure 1, yellow/gold) and the Activation loop (Figure 1, A-loop, red). The A-loop contains a highly conserved Asp-Phe-Gly (DFG) motif that can be either pointing towards the ATP binding site^3^ (DFG in) or rotated away (DFG out). Since all of our present simulations were run in the presence of ATP, we only observed dynamics within the DFG in state. The adenine and ribose rings make hydrogen bonds and van der Waals contacts with the Hinge residues while the phosphate moieties chelate two magnesium ions,^3^ which in turn are ligated by multiple residues within the binding pocket. The N-lobe also contains a conserved helix called the catalytic helix (Figure 1, C-helix, orange). Sequence and structural alignment^25^,^26^ have further highlighted the presence of two non-contiguous structural motifs called the regulatory (R) and catalytic (C) spines, which are required for stabilizing the active state. Sequence alignment has also revealed a conserved glycine-rich loop, or P-loop (Figure 1, green), which connects the β1 and β2 strands^14^,^27^. The loop typically has the Gly-X-Gly-X-X-Gly sequence where X indicates a random amino acid. The P-loop interacts with the phosphate groups on ATP^25^ to position the gamma phosphate for catalysis^28^.

**Figure 1.**
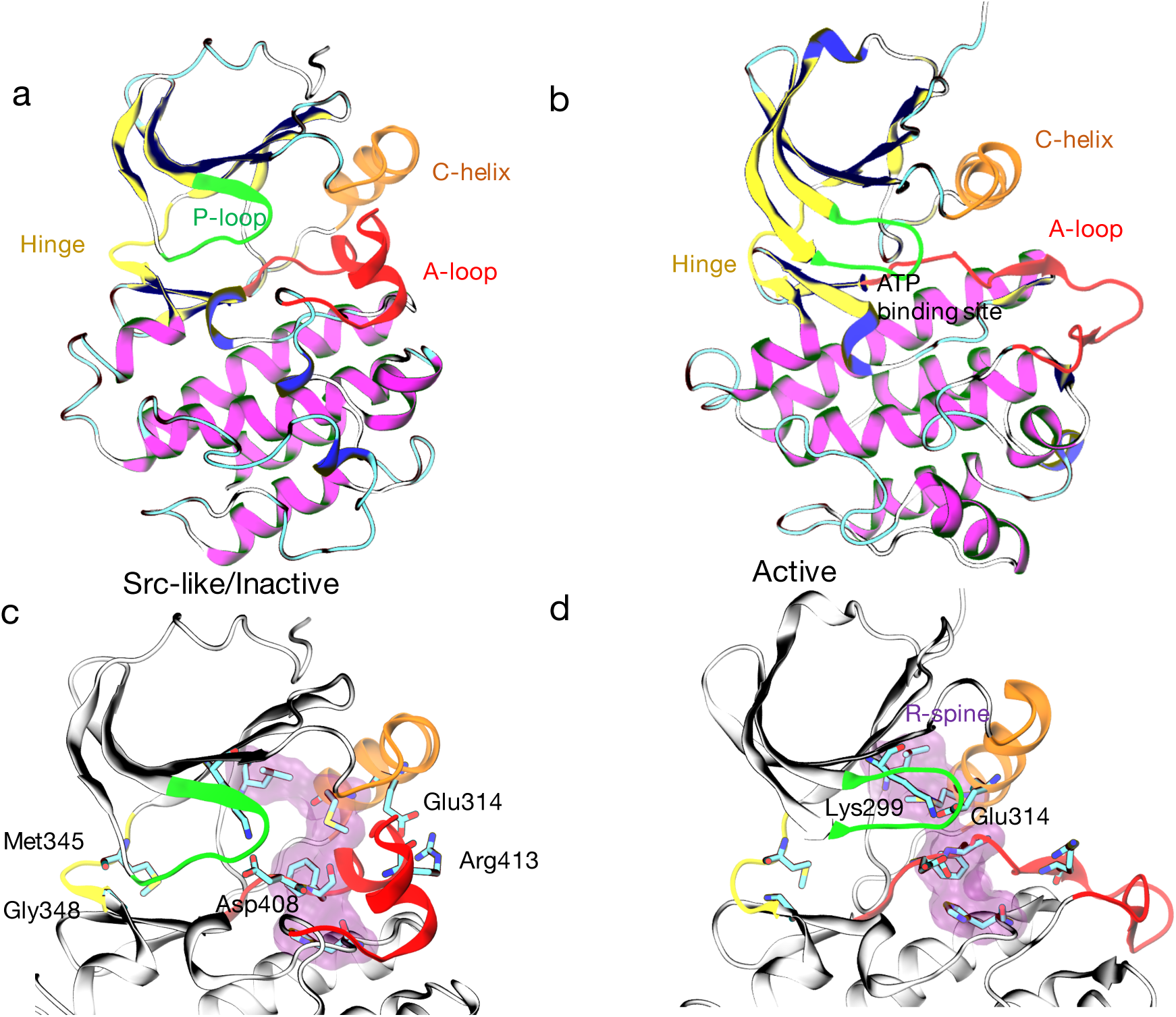
Starting structures for the Fyn simulations were generated from a Src-based homology model corresponding to the inactive state and a crystal structure of Fyn in its active state^22^. In the inactive state (a), the A-loop(red) is folded forming a salt bridge between Glu314 and Arg413. The C-helix (orange) is rotated outwards, and the Hinge (yellow) is open. In the active state (b), the Hinge closes by forming backbone hydrogen bond between Met345 and Gly348, the C-helix rotates inwards to form new contact between Glu314 and Lys299 and complete the hydrophobic R-spine (purple),and the A-loop is unfolded. (c) and (d) show atomistic details of these transitions. ATP and Mg ions have not been rendered for the sake of clarity. Secondary structure was assigned using dictionary of secondary structure in proteins (DSSP)^40^. The color scheme shown here is extended unto the rest of the paper.

While the deactivation pathways for Src^1^,^24^,^29^, and other select kinases^2^,^6^,^18^,^19^ have been extensively studied in the last few years, it remains to be seen whether the deactivation pathways proposed within these prototypical systems can be extended to other kinases. Computational studies of SFK dynamics is challenging due to the prohibitive time-cost of running and analyzing milliseconds of atomistic simulations^30^. Although a few large-scale molecular dynamics (MD) studies^1,15^ have been performed, these have generally relied on biased reaction coordinates for mapping the thermodynamic landscape of kinases^1,31–33^. While these methods could potentially allow for faster converged sampling, there is an inherent risk of biasing the simulation results towards unphysical transition paths^34^. However, given recent work in the development of reaction coordinate free Accelerated Molecular Dynamics (AMD)^35,36^ combined with the ability to run many unbiased parallel simulations on distributed computed platforms^37,38^, we can now reach relevant timescales^39^ to sample the kinase conformational landscape. These parallel trajectories can be stitched together using the Markov state model (MSM) framework to provide additional insight.

Previous MD studies on other kinases^9^,^13^,^16^,^25^,^27^,^32^,^41^,^42^ point to a large conformational change that requires breaking of a conserved Glu-Lys salt bridge, rotation of the C-heliX, folding of the Activation loop (A-loop), and misalignment of regulatory and catalytic spines to form an inactive state. However, the consistency of the transition paths across the kinome and the differences in the accessible energy landscapes for different kinases remain unanswered. Differences in pathways and energy landscapes are central to the rational design of selective type I,II and III inhibitors^2^,^43^ that select for potentially unique intermediate or inactive kinase states rather than the putative active crystal conformation. To answer these questions within the context of the SFKs, we started by sampling the entire deactivation pathway for Fyn starting from a single crystal structure^22^ and a homology model of the inactive catalytic domain for Src^4^, ^13^(see Methods for details). We next used the MD generated Fyn dataset to start thousands of new MD trajectories for six other members of SFK, Lyn, Lck, Hck, Fgr, Yes, and Blk. Ultimately we collected over 5.7ms (Supporting Figure 3) of aggregate MD data on the Folding@home distributed computing platform^37^.The aggregate simulation time was necessary to accurately capture slow conformational transitions (C-helix rotation, A-loop folding), and their relative kinetics (~ 100*s μs*) and thermodynamics across the 7 sequences.

To analyze this data, we created a multiple sequence extension to MSMs^44–47^(see Methods for details), utilizing state-of-the art dimensionality reduction^48^, model selection^49^, and model interpretation^50^ procedures (Supporting Figures 1-2) from Machine Learning (ML) literature. The multi-sequence extension to MSMs allowed us to explicitly compare the thermodynamics and kinetics across the 7 sequences. Our simulations reveal that in the absence of their regulatory domains, the members of the SFK catalytic subunit have active populations ranging from 4-40% and exchange timescales on the order of hundreds of microseconds. We identified several catalytically inactive intermediate states and describe the transition pathways that separate active from inactive states in an effort to gain insight into the atomistic underpinnings of the SFK kinase domain regulatory mechanism. Furthermore, we hope an enhanced picture that connects structural with functional details can set the stage for inhibitor design strategies that i) can take advantage of intermediate states with druggable pockets, and ii) also allow allosteric inhibition and inducible pocket strategies to be conceived. We will begin our paper by first delineating the common activation pathway across the sub-family, focusing on Fyn, before delving into the rest of the sub-family. We are choosing to focus on single sequence because our individual analysis indicated that several members had qualitatively similar activation dynamics.

## Results

### ATP bound Fyn’s samples a kinome wide “DFG in” conformational landscape

In order to quantify the structural heterogeneity within our ATP bound Fyn MD dataset, we compared our results (Figure 2) against the kinome-wide classifications scheme of Möbitz^51^. In that work, the author classified several thousand kinase crystal structures based upon the positioning of conserved DFG (Asp408-Gly410) and C-helix Glu314 residues. Our simulations (Figure 2a) predict that the ATP bound “DFG in” Fyn kinase samples a kinome wide distribution (Figure 2b). However, the “DFG out” (Figure 2 bottom panel) state is inaccessible due to the presence of ATP. This result, combined with our previous work on the apo BTK^42^ catalytic domain, suggests that protein kinases likely possess a conserved conformational landscape, modulated via the exact sequence, post-translational modifications, and the presence of nucleotides, binding partners and substrates. It also shows how MD is increasingly capable of predicting pharmacologically relevant unseen states that have only been previously observed in related members of the super-family.

**Figure 2.**
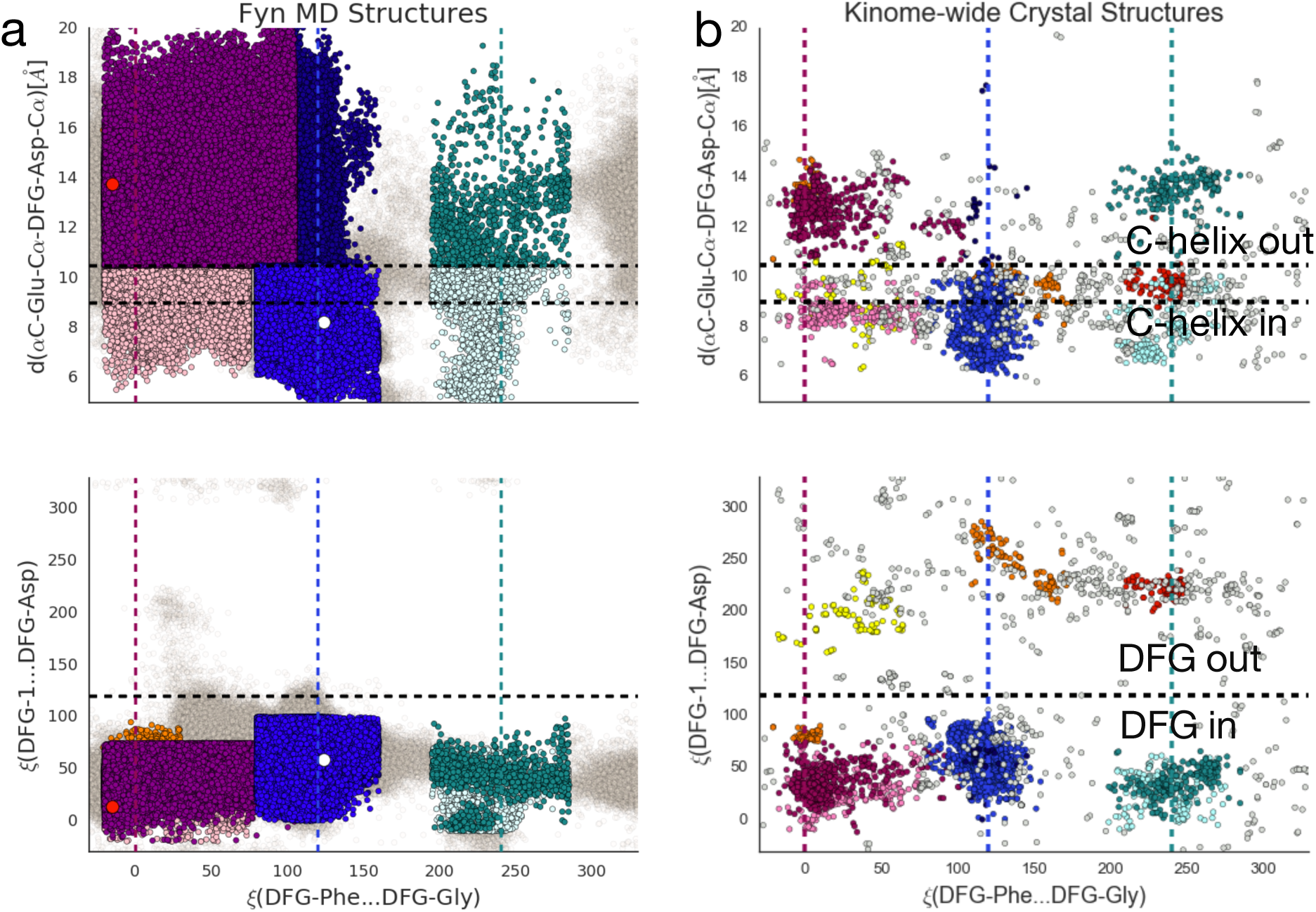
Comparison of Fyn MD dataset, subsampled to 1ns, against the kinome wide conformational diversity reported by Möbitz et al^51^. We used the data and classification scheme provided in ref. ^51^ to generate panel b. The bottom y-coordinate tracks the “DFG in” to “DFG out” transition while the top y coordinate tracks the C-helix in to C-helix out transitions. Note that the “DFG out” state is inaccessible while ATP is bound to the kinase. The x-axis separates the structures along various positions of the DFG motif. The white and red circles represent the starting active (2DQ7) crystal structure and inactive homology model states. For more details regarding the coloring see Supporting Note 1. For the free energies along these coordinates, see Supporting Figure 13.

### Simulations suggest a step-wise activation pathway across the Src sub-family

In order to understand the universality of the kinase activation pathway, we began by simulating the atomistic dynamics of Fyn kinase’s catalytic domain. We connected the active Fyn PDB (RCSB id 2DQ7)^22^ to a homology model^52^ of the Src-like inactive state (Sequence identity 85%) via MD. This required approximately 6 microseconds of accelerated molecular dynamics (AMD) to find seed structures followed by an aggregate 1500 microseconds of sampling on the Folding@home^37^ distributed computing platform. Since neither AMD nor MD requires pre-specifying a reaction coordinate^35^,^53^, we do not expect homology modeling of the inactive state to bias our dynamics. The resulting trajectories were analyzed by time-structure based independent component analysis (tICA)^48^ and MSMs^44^,^45^. tICA is a dimensionality reduction technique designed to find a set of minimally overlapping linear combinations of the input features, such as dihedrals or inter-residue distances, that de-correlate the slowest^48^. The dominant components, called tICs, are one dimensional projections of large scale and slow protein conformational change (Supporting Figure 4). tICA is used as a dimensionality reduction step in MSMs for defining a kinetically motivated clustering metric. The clustering metric defines a state space which is then used as input for the MSM. MSMs are models of dynamic processes defined over a set of protein states and the transition probabilities connecting them^44^,^45^.

Within our Fyn MSM (Figure 3), Hinge closing is highly coupled to activation (Figure 3c, yellow vs. orange trace, respectively). In the Src-like inactive state^27^,^54^,^55^ (Figure 3a, Src-like), the A-loop is folded (Figure 3c, red), the C-helix is rotated outwards (Figure 3c, orange) forming a Glu314-Arg413 salt bridge, and the Hinge is open. The open Hinge is characterized by a broken hydrogen bond (Figure 3c, missing yellow trace) between the backbone carbonyl and amide groups of Met345 and Gly348. Closing of the Hinge (Figure 3c, yellow trace starting at ~90 μs mark), pushes the kinase to an ensemble of intermediate states (Figure 3a, I1) that has a closed/formed Hinge, an unstructured A-loop, and an outward rotated C-helix. Starting from these intermediates, the C-helix can rotate inwards to form a conserved Glu314-Lys299 salt bridge (Figure 3a, Active). This Glu-Arg to Glu-Lys switching mechanism has been observed in active and inactive crystal structures^1^,^13–15^,^33^,^56^ but the intermediate has only been computationally observed^1^,^13^. Within our simulation set, we observed several large-scale activation or deactivation events with several trajectories spontaneously rotating the C-helix in and out (Supporting Figure 5-6).

**Figure 3.**
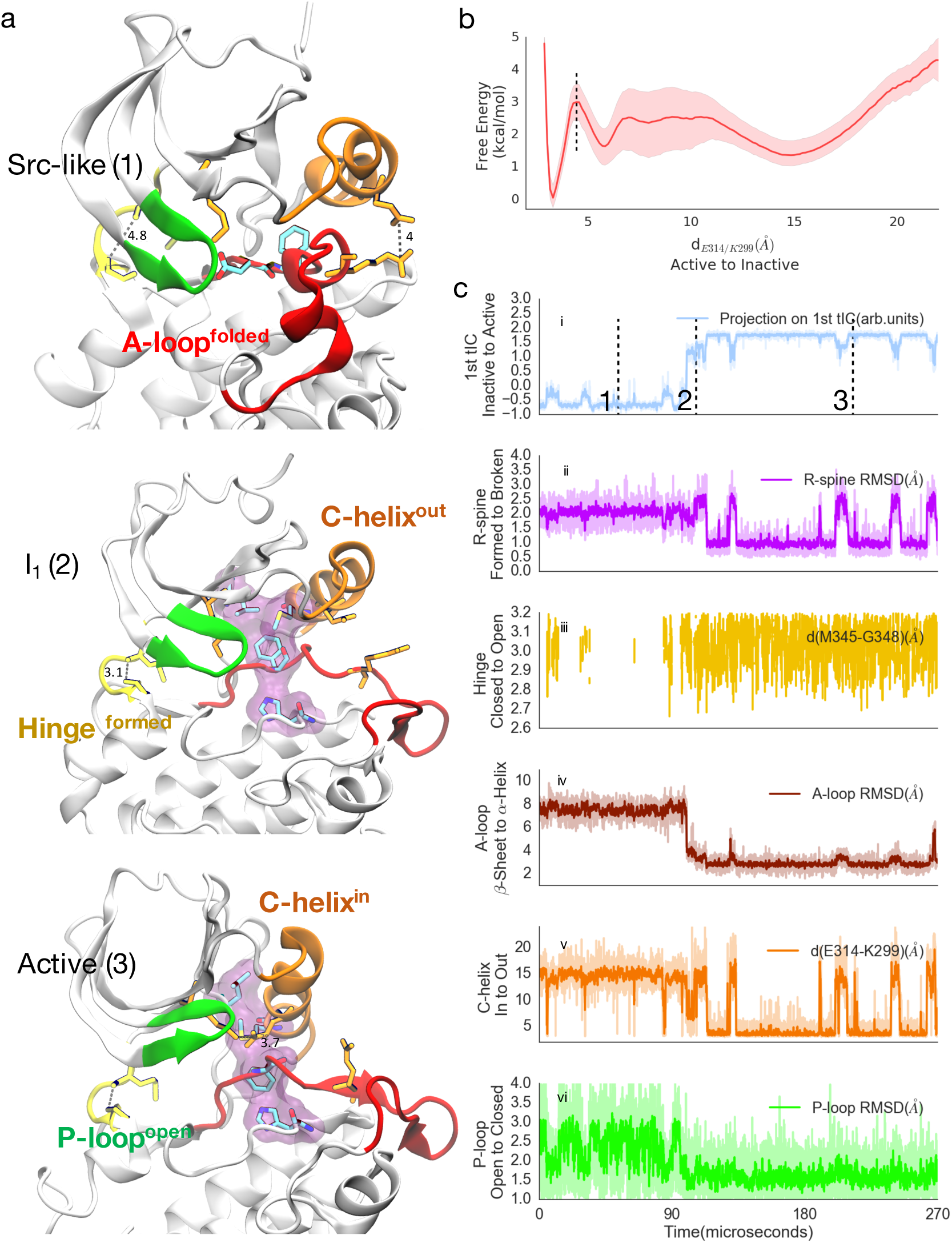
Activation is a concerted process. Model for Fyn kinase starting from the Src-like inactive state (1) and ending in the active state (3). (a) In the Src-like state, the Hinge is broken the kinase is inactive, and the A-loop folds into a double helix. In the Hinge formed states (2-3), the kinase can be active (3) or sample other catalytically inactive intermediates (2). (b). Projection of the Fyn ensemble onto the Glu314-Lys299 distance reaction coordinate gives us the thermodynamics, showing an appreciable active (3) state. The dashed line (Supporting figure 7) indicates the cutoff used to define the active state and the lighter band indicates distributions calculated via 200 rounds of bootstrapping. (c) Kinetics of several molecular switches as inferred by a simulated 270 ***μs*** Monte Carlo MSM trajectory (see methods for details). The traces show variation of several key order parameters. (i) Projection of the trajectory unto the dominant tIC. The 1^st^ tIC captures the Hinge closing and A-loop dynamics. The black dashed line corresponds to the structures plotted in (a). (ii) The second panel shows the R-spine RMSD. (iii) The third panel shows the hydrogen bond distance between the backbone atoms of Hinge residues M345 and G348. Missing values indicate the absence of a hydrogen bond, calculated according to the Baker Hubbard criterion^57^. (iv) The fourth panel traces the RMSD of A-loop heavy atoms (residue Lys405-Gln424). (v) The fifth panel shows the distance between residues Glu314 (Delta Carbon) and Lys299 (Zeta Nitrogen). Larger values indicate outward swing of the C-helix. (vi) The last panel shows the RMSD of the P-loop. All RMSDs are calculated using the heavy atoms and are in reference to the active structure. The colors correspond to the structural features labeled in Figure 1 and darker colors in (c) indicate moving averages across 10 frames.

In particular, the intermediate state (Figure 3a I1, Supporting Figure 7) observed within our ensemble is different from either of the starting active (2DQ7) and inactive homology model states. In this state, the Hinge is closed, the C-helix is rotated outwards but the A-loop is relatively unstructured and samples a range of conformations^1^. The outward rotation of the C-helix opens up a pocket between the ATP binding site and the C-helix (Supporting Figure 10) that presents itself as a putatively druggable site for the design novel allosteric Fyn inhibitors (Supporting Figure 10) – not too dissimilar from previous observations that were made in the context of Src^1^.

The activation process is accompanied by the alignment of two hydrophobic spines^25^,^26^, the Catalytic spine (C-spine; Val285, Ala297, Leu397, Ile396, Val398, Leu350, Leu455, Leu459, and ATP) and the Regulatory spine (R-spine; Leu329, Met318, Phe409, His388). These noncontiguous structural motifs are aligned in the kinase active state. The C-spine is broken by the movement of ATP’s adenine and ribose rings towards the protein core^15^ while the R-spine is disrupted by movement of the C-helix away from the core (Figure 1), leading to a displacement of Met318 and Phe409. Lastly, activation correlates with a change in positioning of the glycine rich P-loop (Figure 3c, green trace). Though both ensembles occupy the opposing configurations, our simulations find that the inactive ensemble favors a closed P-loop configuration whilst the active ensemble favors the open (Figure 3c) state.

### Active and inactive states are similarly populated

In order to gain insight into Fyn’s thermodynamics, we projected the Fyn ensemble onto a simplified reaction coordinate (Figure 3b, Supporting Figure 8) measuring the contact distance between the SFK conserved C-helix Glu314 and catalytic Lys299 residues (numbering corresponds to Fyn). Our analysis indicates that the active state is similar in population to the inactive state (Figure 3) in the Fyn ensemble 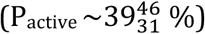. The sub and superscript represent 25^th^ and 75^th^ percentiles as calculated via 200 rounds of bootstrapping (see Methods for details, and Figure 4 and Supporting Figure 9 for the complete distribution). The active state population within our model is consistent with the hypothesis that the kinase regulatory domains (SH2 and SH3) are required to regulate SFK^13,27^ kinases, and their removal leads to a higher active population. The appreciable active population (Figure 3) within our model also helps to potentially explain the experimental phosphorylation patterns of SFKs. SFKs require phosphorylation of a key tyrosine residue (Tyr420) in the activation loop (A-loop) for activation^27^,^55^. While we were unable to find phosphorylation data for Fyn for a direct comparison to our model, phosphorylation was shown to have a minimal effect upon Src, featuring an increase in activity of only about 1.5-2.5x^58–60^. In contrast, phosphorylation increases the active population in non-SFK extracellular signal regulated kinase 2 (ERK2) by three orders of magnitude^61^. We predict Fyn to behave similarly to Src given their 85% sequence identity and the results of our un-phosphorylated simulations. This increase in activity is consistent with a thermodynamic landscape featuring an appreciably active minimum that is robust to phosphorylation (characterized by population shifts of less than an order of magnitude or 1 kcal/mol). These small shifts in active population are expected to result in relatively minor increases in observed chemical activity.

To gain insight into Fyn’s activation kinetics, we calculated the distribution for the median time required to “travel” to a set of active states (Figure 3, Supporting Figure 7) starting from the Src-like inactive states (Supporting Figure 7). According to our model, the median value for Fyn’s mean first passage time to either of its active or inactive states is on the hundreds of microsecond timescale (Figure 4b, Supporting Figure 9). Such exceptionally high timescales make it difficult, even with millisecond scale simulations, to provide reliable kinetic estimates. Accordingly, we can only conservatively estimate that the exchange timescales are on the micro to millisecond timescale.

Similar to previous experimental work on protein kinase A^62^, ERK2^61^, p38^62^, and Src^63^, the concerted conformational change predicted by our simulations is likely to display NMR signature. In particular, our MSM is highly consistent with the Src study^63^ where the authors experimentally showed that Src’s catalytic domain exists in at least two conformational states in solution. The major and minor conformational state exchange on the micro to millisecond timescales and are related via hinge motion. Similar to their experimental data, our simulations also predict a two state hinge behavior with the hinge formed states having a higher free energy (<1-2kcal/mol) relative to the hinge broken states. The hinge formed states can be further divided into the active and intermediate. We predict that similar experiments on Fyn can be used to probe the multi-state behavior for the the salient features of our activation model, including allosteric coupling of the hinge residues to the rest of the kinase.

### High sequence identity leads to similar in-silico functional dynamics

In order to build a single model for Src sub-family, we ran new simulations for several SFK members. To that end, we extracted several hundred structures from our Fyn simulation set, built homology models of the human sequences for six SFK members (Lyn, Lck, Hck, Fgr, Yes, and Blk), and docked ATP + magnesium ions into each ATP binding site (Figure 1). This large pool of configurations allowed us to seed simulations from high and low free energy regions in the kinase state space, enabling faster convergence of sampling^44^,^64^. We furthermore validated the active and inactive homology models by comparing them to known crystallographic states (Supporting Figure 11). Based on their high sequence similarity with Fyn (Supporting Note 1, Figure 4a), we conjectured that these systems would be expected to display similar dynamics.

A set of MD simulations was carried out for each of these kinase systems, resulting in six sets of trajectories with an aggregate sampling of 4 ms. A separate MSM was built for each of the six kinase domains, utilizing an identical set of clustering definition. This single domain decomposition allowed us to explicitly compare the thermodynamics and kinetics of Fyn, Lyn, Lck, Hck, Fgr, Yes, and Blk, the results of which are distilled into Figure 4.

Collectively, within the 1kcal/mol restraints of modern force fields, our simulations suggest that the free energy differences between active and inactive ensembles fall within 1-2kcal/mol across all sequences (Figure 4a and Supporting figure 7-8), with active state populations ranging from 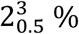 for Lyn to 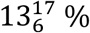 for Hck to 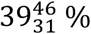 for Fyn. As before, the sub- and superscript represent 25^th^ and 75^th^ percentiles as calculated via 200 rounds of bootstrapping (see Figure 4 and Supporting Figure 9 for the complete distribution). The active states of both Fyn and Hck kinase domains are characterized by elevated populations, which is qualitatively consistent with experimental evidence that both full-length kinases display higher specific activities when compared to Lyn^65^.

**Figure 4.**
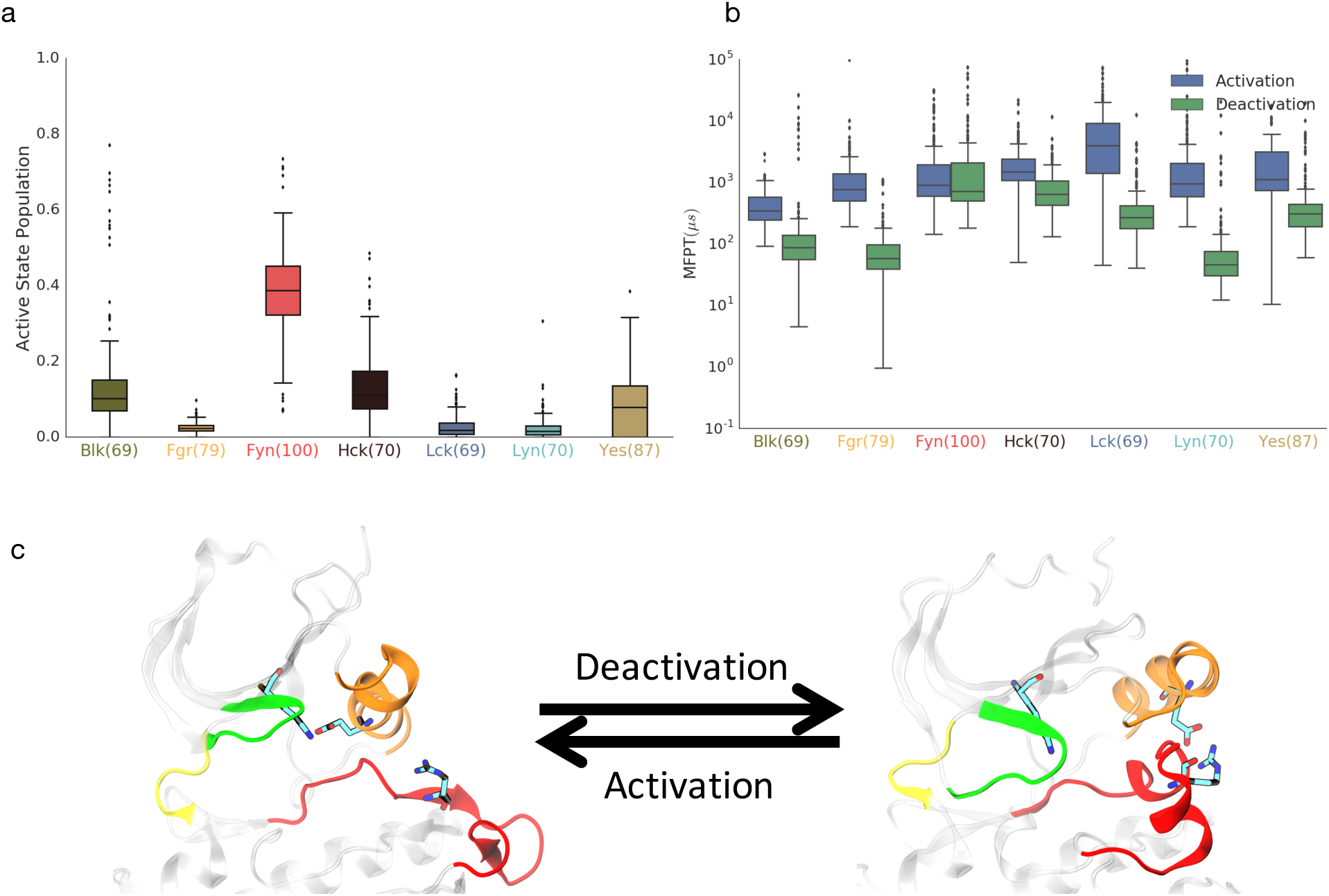
The catalytic domains of SFK members sample several macrostates with active states being within 2kcal/mol of the inactive state. (a) Boxplot distributions for the active state populations for all 7 proteins. We used a consistent MSM basis to allow for direct comparisons across the sequences. See Supporting Figure 8, for projections along the activation coordinate. The bracketed numbers correspond to sequence identity to Fyn. (b) Boxplot distribution for activation and deactivation bootstrapped mean first passage time (MFPT) for all 7 proteins. We used the same set of starting and ending states for the calculations, though not all states are accessible for the varying sequences. The box plot box shows the 25th and 75th quartiles while the whiskers show the rest of the distribution. Outliers (defined as being 1.5x outside the quartile values) are shown as individual points. All timescales > 10**5 (0.26%) were discarded. (c) Frames showing representative active and inactive states used for the calculation in 3a and 3b. See Supporting Figure 7 for the exact definition of the active and inactive states, and Supporting Figure 9 for the future statistical validation. ATP and Mg have not been rendered for sake of clarity.

A kinetics analysis for all six kinases (Figure 4b) shows that similar to Fyn, all six SFK members display activation timescales (Figure 4b, blue boxes) on the order of milliseconds and deactivation timescales (Figure 4b, green boxes) on the order of 100-1000 μs. While the distributions show overlap, for most of the simulated sequences, deactivation was faster than activation except for Fyn and Hck (Figure 4b). The asymmetric timescales are consistent with the presence of a slightly higher free-energy active state (Figure 4a). The higher free energy state indicates that phosphorylation might be required for complete kinase activation. The slower Fyn and Hck deactivation is consistent with their higher observed experimental activities. Therefore, while we can predict that SFK will exchange on the hundred microsecond to millisecond timescale, more precise kinetic predictions will likely require simulations on the order of tens to hundreds of milliseconds.

## Conclusions

Unbiased, multi-millisecond MD dynamics of seven SFK members presented here have provided us with detailed atomistic insight into their collective activation mechanisms, revealing the prominence of Hinge motion for activation. Our simulations indicate that SFK sample a conserved free energy landscape which has been previously only observed at the kinome level^51,66^. Furthermore, the activation process further correlates with dampening of dynamics within the P-loop, in addition to large scale conformational changes involving the C-helix and A-loop. Our Fyn MSM suggests that the catalytic domain, in the absence of its regulatory partners, has a measureable active state and contains several catalytically inactive intermediate and inactive states.

We present an extension to traditional MSMs built on top of a set of sequence invariant feature space. The joint phase space-decomposition allows for a comparative analysis of the kinetics and thermodynamics across the entire sub-family. Our calculations predict that the active state is within 1-2 kcal/mol of the inactive ensemble, and that activation is slower than deactivation across the entire sub-family. The transitions between active and inactive states proceed through a number of intermediate conformations that offer new opportunities for the design of kinase subfamily specific inhibitors.

While we have chosen to look at the conserved dynamics within the current dataset, an interesting complementary extension might involve looking at the dynamics of the nonconserved residues in an effort to elucidate differences within the SFK family. More intriguingly, our methodology raises opportunities for creating a single kinome-wide model, allowing researches to compare and contrast the thermodynamic and kinetic effects of sequence changes, oncogenic mutations, and PTMs to a given baseline. Such unbiased comparative structural modeling can aid in the creation of new, specific and potent kinase inhibitors or perhaps even creating personalized medicines conditioned upon the patients’ genome.

## Methods

### Initial Setup and homology modeling

The crystallographic coordinates of Fyn kinase were downloaded from the protein databank (id: 2DQ7)^22^. The staurosporine drug and crystallographic waters were deleted from the pdb file leaving just the protein coordinates. The structure was mutated to the human WT sequence using Modeller (ver. 9.1). VMD (ver. 1.9)^67^ was used to add adenosine tri phosphate(ATP) to the structures using the Src^1^ structure as the template. Modeller^52^ was further used to build the Fyn’s inactive structure (with ATP) using the inactive c-Src structure(id: 2SRC)^1^,^13^,^16^ as a template. Hydrogen atoms were added to both the active and inactive structure using the t-leap module contained with the Amber tools (ver. 14) suit.^68–^ ^70^ Both the systems were solvated in a water box with 10 Å padding on all side. Chloride and sodium ions were added to neutralize the charge and set up the final ionic concentration to 150mM. The Amber99sb-ildn^71^ force field was used to model protein dynamics in conjunction with the TIP3P^72^ water model and Amber ATP and Mg parameters^73^.

### Simulation Details

All production runs for the initial accelerated dynamics^35^ and regular molecular dynamics simulations were run at a pressure of 1 atm and temperature of 300K. The Langevin integrator with a friction coefficient of 1/ps and a 2fs time step were used. A Monte Carlo barostat with an interval of 25 frames was used to maintain the pressure at 1ATM. Frames were saved every 100ps unless otherwise specified. Long-range electrostatics were dealt with using the Particle mesh Ewald^74^ algorithm with a 8Å cutoff for the accelerated molecular dynamics simulation and a 10Å cutoff for the production runs on Folding@home^37^. All simulations were equilibrated for 1ns before production runs unless specified otherwise.

### Accelerated Molecular Dynamics Simulations

To obtain a large number of diverse initial starting structures, accelerated molecular dynamics (AMD)^35^ was run on both the active and the inactive starting structures until partial deactivation and activation were observed respectively (~6 μs) as monitored via progress on the two dimensional reaction coordinate defined using the partial unfolding of the A-loop and swinging of the C-helix. For the accelerated dynamics, a boost was applied to the dihedrals of the system with an energy threshold of 3900-4500kcal/mol and an alpha parameter value of 160-210. 1005 structures were randomly pulled from the resulting trajectories to seed simulation on the Folding@Home^37^ distributed computing platform.

### Production MD Simulations

The 1005 randomly pulled structures from the AMD trajectories were re-solvated and minimized as described above using the Amber Tools suite^68^. The resulting Amber topology and input coordinates were read into OpenMM(ver. 6.3-7.1)^75^ for production runs on FAH. 1.53ms of combined dynamics were obtained in three stages (project id 9101-9102, 9162) consisting of approximately equal sampling. The later stages involved sampling from the lowly populated states.

### SFK homology modeling and Production MD simulation

For Lyn, Lck, Hck, Fgr, Yes, and Blk kinase simulations, we sampled 1000 structures from the Fyn kinase dataset and homology modeled the human sequence onto those structures. ATP, Mg ions, water and counter ions were added as before. We used the same protein force field and water models, and equilibrated the models using the protocol described above. For production MD simulations, we again employed a similar protocol but saved trajectory frames every 200ps to save disk space.

### Markov state model

Before model building, all 7 datasets were subsampled to 1ns to allow for faster analysis. Building an MSM requires identification of metastable kinetically close conformational states. This splitting of the accessible phase space is followed by counting the transitions between those states as observed in our trajectories at a Markovian (memory free) lag time. After sampling, a total of 13343 trajectories (across 7 sequences) were vectorized using a common set of 1081 protein contacts. This was done by calculating all pair wise contacts for residues W264, E265, G280, F282, G283, V285, W286, V296, K299, M306, F311, L312, E314, A315, M318, K319, L321, L326, L329, A331, E336, H388, R389, D390, L391, D408, F409, G410, L411, A412, R413, I415, D417, E419, Y420, G425, K427, F428, K431, E436, G441, K446, S451, T461, P466, C491, and P511. The residue numbering here corresponds to Fyn’s and we used equivalent contact distances across all proteins. The Supporting Information contains a sequence alignment that was used for this purpose. Our method is completely generic and can be extended to many proteins. For new set of systems, we recommend first performing a sequence alignment and then using a conserved set of residues to define the feature basis. Care must be taken if some sequences have insertions or deletions since that would preclude the use of certain backbone dihedrals as features. We note that the open source software MSMBuilder implements a range of utility functions that make this task easier.

The featurized trajectories were then transformed using time structure independent component analysis (tICA)^48^. tICA seeks to find a set of linear combinations of features that de-correlate the slowest (at a certain lag time) whilst minimizing their overlap. The transformed dataset was clustered using the mini batch K-means. It is worth noting that we built a single tICA and K-Means model for all 7 sequences allowing us to quantitatively compare thermodynamics and kinetics. We use a tICA lagtime of 300ns to reduce the dimensionality of the data from 1081 contacts to 5tICs, which were also scaled using kinetic mapping^76^. For the clustering step, we used the mini batch K-Means algorithm with the number of clusters set to 300. Based upon previous work^1^,^38^ and the convergence of the implied timescales plot for 300 state model, we chose a lag time of 30ns (Supporting Figure 1) for the MSM. The Markov transition matrix was fit via Maximum likelihood estimation (MLE) with reversibility and ergodicity constraints. This procedure discarded less than 1% of the data for any given sequence indicating converged sampling. See Supporting Figure 1 and Figure 9 for further validation.

### Error analysis

To obtain error bars for the MFPTs, timescales, and equilibrium populations, 200 rounds of bootstrapping were performed for each simulated protein sequence. The bootstrapping was performed over the trajectories so that for each bootstrap sample, we randomly picked N trajectories (with replacement) where N is the total number of trajectories for that sequence. We kept the final state labeling, but fit a *new* MSM to this resampled trajectory set. This leads to a new estimate of the thermodynamics and kinetics. Repeating this 200 times allows us to estimate error bars for the predicted kinetics and thermodynamics. We also validated our results by restricting each trajectory to between 75 and 100% of its final length (Supporting Figure 9) and re-computing all reported thermodynamic and kinetic statistics and found them to be consistent.

The trajectories were featurized and analyzed using the MDTraj (ver. 1.7-v1.8)^77^ package while tICA dimensionality reduction and Markov modeling were performed using MSMbuilder (ver. 3.5+)^78^. Most of the analysis was performed within the IPython scientific environment^79^ with extensive use of the MSMExplorer, matplotlib (ver. 1.0+)^80^, and scikit-learn libraries^81^. All protein images were generated using visual molecular dynamics (VMD, ver. 1.9+)^67^ and all protein surfaces were rendered using SURF^82^ as implemented in VMD.

### Model interpretation

The models were primarily analyzed using techniques laid out in previous papers^83^,^50^. To further query the model, we inferred the dominant deactivation pathways using Transition path theory (TPT) ^84^,^85^. TPT requires specifying the starting and ending states to find the interconnecting pathways. All states with a formed Hinge(1^st^ tIC value > 1.5) and Glu314-Lys299 bridge (< 4.45Å) were considered active. For the inactive states, we picked all states whose all heavy atom RMSD was with 4 Å of the homology modeled Src-like inactive state. The intermediate state was defined to have intermediate values (between 0.5 and 1.3.) along the first tIC.

We also queried the model by generating long trajectories from the MSM. Starting from the inactive state, we propagated the dynamics at a lag time of 30ns using a random number generator and the Markovian transition model as our sampling scheme. At each “timestep”, we randomly picked a frame assigned to that state to report the instantaneous observable. Given correct parameterization of the underlying transition matrix, this method produces dynamics comparable to running a single long MD trajectory.

The transition pathways were analyzed using a combination of inspection using visual molecular dynamics (VMD)^67^, random forest classifiers^50^, and tICA projections^48^,^83^. The slowly de-correlating tICA vectors were used to find the degrees of freedom involved in the dynamics^83^. Spectral decomposition of the MSM transition matrix was used to estimate the equilibrium populations and dynamical processes connecting those Markov states. The second eigenvector corresponding to the slowest dynamical process correlates very highly with the active to inactive process (Supporting Figure 2) with an associated timescale of 100μs (Supporting Figure 1), consistent with previous theoretical work^1^.

## Acknowledgments

We thank the donors of the Folding@home distributed computing platform for graciously providing their compute power used for this project. M.M.S would like to acknowledge support from the National Science Foundation grant NSF-MCB-0954714 and the NIH S10 Shared Instrumentation Grant 1S10RR02664701 for their support of the Biox3 computer cluster at Stanford. GK acknowledges support from the the NIH Simbios Program, and the Center for Molecular Analysis and Design at Stanford. We would also like to thank Diwakar Shukla, Matthew Harrigan, Brooke Husic, Ariana Peck, Jade Shi, Carlos Hernández and other members of the Pande Lab for many insightful discussions and critical comments on the manuscript.

## References

1. Shukla, D., Meng, Y., Roux, B. & Pande, V. S. Activation pathway of Src kinase reveals intermediate states as targets for drug design. Nat. Commun. 5, 3397 (2014).

2. Skora, L., Mestan, J., Fabbro, D., Jahnke, W. & Grzesiek, S. NMR reveals the allosteric opening and closing of Abelson tyrosine kinase by ATP-site and myristoyl pocket inhibitors. Proc. Natl. Acad. Sci. U. S. A. (2013). doi:10.1073/pnas.1314712110

3. Endicott, J. A., Noble, M. E. M. & Johnson, L. N. The structural basis for control of eukaryotic protein kinases. Annu. Rev. Biochem. 81, 587–613 (2012).

4. Johnson, L. N., Noble, M. E. M. & Owen, D. J. Active and inactive protein kinases: Structural basis for regulation. Cell 85, 149–158 (1996).

5. Manning, G., Whyte, D. B., Martinez, R., Hunter, T. & Sudarsanam, S. The protein kinase complement of the human genome. Science 298, 1912–34 (2002).

6. Nagar, B. et al. Structural basis for the autoinhibition of c-Abl tyrosine kinase. Cell 112, 859–71 (2003).

7. Nowakowski, J. et al. Structures of the cancer-related Aurora-A, FAK, and EphA2 protein kinases from nanovolume crystallography. Structure 10, 1659–67 (2002).

8. Greuber, E. K., Smith-Pearson, P., Wang, J. & Pendergast, A. M. Role of ABL family kinases in cancer: from leukaemia to solid tumours. Nat. Rev. Cancer 13, 559–71 (2013).

9. Parsons, S. J. & Parsons, J. T. Src family kinases, key regulators of signal transduction. Oncogene 23, 7906–9 (2004).

10. de Wispelaere, M., LaCroix, A. J. & Yang, P. L. The small molecules AZD0530 and dasatinib inhibit dengue virus RNA replication via Fyn kinase. J. Virol. 87, 7367–81 (2013).

11. Schenone, S. et al. Fyn kinase in brain diseases and cancer: the search for inhibitors. Curr. Med. Chem. 18, 2921–42 (2011).

12. Akimoto, M. et al. Signaling through dynamic linkers as revealed by PKA. Proc. Natl. Acad. Sci. 110, 14231–6 (2013).

13. Xu, W., Harrison, S. C. & Eck, M. J. Three-dimensional structure of the tyrosine kinase c-Src. Nature 385, 595–602 (1997).

14. Cowan-Jacob, S. W. et al. The crystal structure of a c-Src complex in an active conformation suggests possible steps in c-Src activation. Structure 13, 861–71 (2005).

15. Foda, Z. H., Shan, Y., Kim, E. T., Shaw, D. E. & Seeliger, M. A. A dynamically coupled allosteric network underlies binding cooperativity in Src kinase. Nat. Commun. 6, 1–10 (2015).

16. Bernadó, P., PéreZ, Y., Svergun, D. I. & Pons, M. Structural characterization of the active and inactive states of Src kinase in solution by small-angle X-ray scattering. J. Mol. Biol. 376, 492–505 (2008).

17. Hantschel, O. & Superti-Furga, G. Regulation of the c-Abl and Bcr-Abl tyrosine kinases. Nat. Rev. Mol. Cell Biol. 5, 33–44 (2004).

18. Pendergast, A. M. The Abl family kinases: Mechanisms of regulation and signaling. Advances in Cancer Research 85, 51–100 (2002).

19. Shan, Y., Arkhipov, A., Kim, E. T., Pan, A. C. & Shaw, D. E. Transitions to catalytically inactive conformations in EGFR kinase. Proc. Natl. Acad. Sci. U. S. A. 110, 7270–5 (2013).

20. Xu, W., Doshi, A., Lei, M., Eck, M. J. & Harrison, S. C. Crystal structures of c-Src reveal features of its autoinhibitory mechanism. Mol. Cell 3, 629–38 (1999).

21. Resh, M. D. Fyn, a Src family tyrosine kinase. Int. J. Biochem. Cell Biol. 30, 1159–62 (1998).

22. Kinoshita, T., Matsubara, M., Ishiguro, H., Okita, K. & Tada, T. Structure of human Fyn kinase domain complexed with staurosporine. Biochem. Biophys. Res. Commun. 346, 840–4 (2006).

23. Williams, J. C., Wierenga, R. K. & Saraste, M. Insights into Src kinase functions: structural comparisons. Trends Biochem. Sci. 23, 179–84 (1998).

24. Foda, Z. H., Shan, Y., Kim, E. T., Shaw, D. E. & Seeliger, M. A. A dynamically coupled allosteric network underlies binding cooperativity in Src kinase. Nat. Commun. 6, 5939 (2015).

25. Kornev, A. P., Haste, N. M., Taylor, S. S. & Eyck, L. F. T. Surface comparison of active and inactive protein kinases identifies a conserved activation mechanism. Proc. Natl. Acad. Sci. U. S. A. 103, 17783–8 (2006).

26. Taylor, S. S. & Kornev, A. P. Protein Kinases: Evolution of Dynamic Regulatory Proteins. Trends Biochem. Sci. 36, 65–77 (2011).

27. Sicheri, F. & Kuriyan, J. Structures of Src-family tyrosine kinases. Curr. Opin. Struct. Biol. 7, 777–85 (1997).

28. Shaltiel, S., Cox, S. & Taylor, S. S. Conserved water molecules contribute to the extensive network of interactions at the active site of protein kinase A. Proc. Natl. Acad. Sci. U. S. A. 95, 484–91 (1998).

29. Yang, S., Banavali, N. K. & Roux, B. Mapping the conformational transition in Src activation by cumulating the information from multiple molecular dynamics trajectories. Proc. Natl. Acad. Sci. U. S. A. 106, 3776–3781 (2009).

30. Bowman, G. R., Pande, V. S. & Noé, F. An Introduction to Markov State Models and Their Application to Long Timescale Molecular Simulation. 797, (2014).

31. Pan, A. C., Sezer, D. & Roux, B. Finding transition pathways using the string method with swarms of trajectories. J. Phys. Chem. B 112, 3432–40 (2008).

32. Yang, S. & Roux, B. Src kinase conformational activation: thermodynamics, pathways, and mechanisms. PLoS Comput. Biol. 4, e1000047 (2008).

33. Ozkirimli, E., Yadav, S., Miller, W. & Post, C. An electrostatic network and long-range regulation of Src kinases. Protein Sci. 295, 1871–1880 (2008).

34. Pan, A. C., Weinreich, T. M., Shan, Y., Scarpazza, D. P. & Shaw, D. E. Assessing the accuracy of two enhanced sampling methods using egfr kinase transition pathways: The influence of collective variable choice. J. Chem. Theory Comput. 10, 2860–2865 (2014).

35. Hamelberg, D., Mongan, J. & McCammon, J. A. Accelerated molecular dynamics: a promising and efficient simulation method for biomolecules. J. Chem. Phys. 120, 11919–29 (2004).

36. Abrams, C. & Bussi, G. Enhanced Sampling in Molecular Dynamics Using Metadynamics, Replica-Exchange, and Temperature-Acceleration. Entropy 16, 163–199 (2013).

37. Shirts, M. & Pande, V. S. Screen Savers of the World Unite! Science 290, 1903–1904 (2000).

38. Kohlhoff, K. J. et al. Cloud-based simulations on Google Exacycle reveal ligand modulation of GPCR activation pathways. Nat. Chem. 6, 15–21 (2014).

39. Lane, T. J., Shukla, D., Beauchamp, K. A. & Pande, V. S. To milliseconds and beyond: challenges in the simulation of protein folding. Curr. Opin. Struct. Biol. 23, 58–65 (2013).

40. Kabsch, W. & Sander, C. Dictionary of protein secondary structure: pattern recognition of hydrogen-bonded and geometrical features. Biopolymers 22, 2577–637 (1983).

41. Levinson, N. M., Seeliger, M. A., Cole, P. A. & Kuriyan, J. Structural basis for the recognition of c-Src by its inactivator Csk. Cell 134, 124–34 (2008).

42. Sultan, M. M., Denny, R. A., Unwalla, R., Lovering, F. & Pande, V. S. Millisecond Dynamics Of BTK Reveal Kinome-Wide Conformational Plasticity Within The Apo Kinase Domain. bioRxiv 135913 (2017). doi:10.1101/135913

43. Kuglstatter, A. et al. Insights into the conformational flexibility of Bruton’s tyrosine kinase from multiple ligand complex structures. Protein Sci. 20, 428–436 (2011).

44. Pande, V. S., Beauchamp, K. & Bowman, G. R. Everything you wanted to know about Markov State Models but were afraid to ask. Methods 52, 99–105 (2010).

45. Prinz, J.-H. et al. Markov models of molecular kinetics: generation and validation. J. Chem. Phys. 134, 174105 (2011).

46. Voelz, V. A., Elman, B., Razavi, A. M. & Zhou, G. Surprisal metrics for quantifying perturbed conformational dynamics in Markov state models. J. Chem. Theory Comput. 10, 5716–5728 (2014).

47. Wan, H., Zhou, G. & Voelz, V. A. A Maximum-Caliber Approach to Predicting Perturbed Folding Kinetics Due to Mutations. J. Chem. Theory Comput. 12, 5768–5776 (2016).

48. Schwantes, C. R. & Pande, V. S. Improvements in Markov State Model Construction Reveal Many Non-Native Interactions in the Folding of NTL9. J. Chem. Theory Comput. 9, 2000–2009 (2013).

49. McGibbon, R. T. & Pande, V. S. Variational cross-validation of slow dynamical modes in molecular kinetics. J. Chem. phyics 142, (2015).

50. Sultan, M. M., Kiss, G., Shukla, D. & Pande, V. S. Automatic Selection of Order Parameters in the Analysis of Large Scale Molecular Dynamics Simulations. J. Chem. Theory Comput. 10, 5217–5223 (2014).

51. Möbitz, H. The ABC of protein kinase conformations. Biochim. Biophys. Acta - Proteins Proteomics 1854, 1555–1566 (2015).

52. Eswar, N. et al. Comparative protein structure modeling using MODELLER. Curr. Protoc. Protein Sci. Chapter 2, Unit 2.9 (2007).

53. Dror, R. O., Dirks, R. M., Grossman, J. P., Xu, H. & Shaw, D. E. Biomolecular simulation: a computational microscope for molecular biology. Annu. Rev. Biophys. 41, 429–52 (2012).

54. Levinson, N. M. et al. A Src-like inactive conformation in the abl tyrosine kinase domain. PLoS Biol. 4, e144 (2006).

55. Huse, M. & Kuriyan, J. The conformational plasticity of protein kinases. Cell 109, 275–282 (2002).

56. Huang, H., Zhao, R., Dickson, B. M., Skeel, R. D. & Post, C. B. αC helix as a switch in the conformational transition of Src/CDK-like kinase domains. J. Phys. Chem. B 116, 4465–75 (2012).

57. Baker, E. N. & Hubbard, R. E. Hydrogen bonding in globular proteins. Prog. Biophys. Mol. Biol. 44, 97–179 (1984).

58. Sun, G. et al. Effect of autophosphorylation on the catalytic and regulatory properties of protein tyrosine kinase Src. Arch. Biochem. Biophys. 397, 11–17 (2002).

59. Weijland, A. et al. The purification and characterization of the catalytic domain of Src expressed in Schizosaccharomyces pombe. Comparison of unphosphorylated and tyrosine phosphorylated species. Eur. J. Biochem. 240, 756–764 (1996).

60. Kemble, D. J., Wang, Y. H. & Sun, G. Bacterial expression and characterization of catalytic loop mutants of Src protein tyrosine kinase. Biochemistry 45, 14749–14754 (2006).

61. Xiao, Y. et al. Phosphorylation releases constraints to domain motion in ERK2. Proc. Natl. Acad. Sci. U. S. A. 111, 2506–11 (2014).

62. Xiao, Y., Liddle, J. C., Pardi, A. & Ahn, N. G. Dynamics of protein kinases: Insights from nuclear magnetic resonance. Acc. Chem. Res. 48, 1106–1114 (2015).

63. Pucheta-Martínez, E. et al. An Allosteric Cross-Talk Between the Activation Loop and the ATP Binding Site Regulates the Activation of Src Kinase. Sci. Rep. 6, 24235 (2016).

64. Chodera, J. D. & Noé, F. Markov state models of biomolecular conformational dynamics. Current Opinion in Structural Biology 25, 135–144 (2014).

65. Moroco, J. A. et al. Differential sensitivity of Src-family kinases to activation by SH3 domain displacement. PLoS One 9, e105629 (2014).

66. Sultan, M. M., Denny, R. A., Unwalla, R., Lovering, F. & Pande, V. S. Millisecond dynamics of BTK reveal kinome-wide conformational plasticity within the apo kinase domain. Sci. Rep. 7, 15604 (2017).

67. Humphrey, W., Dalke, A. & Schulten, K. VMD: visual molecular dynamics. J. Mol. Graph. 14, 33–8, 27-8 (1996).

68. Salomon-Ferrer, R., Case, D. A. & Walker, R. C. An overview of the Amber biomolecular simulation package. Wiley Interdiscip. Rev. Comput. Mol. Sci. 3, 198–210 (2013).

69. Götz, A. W. et al. Routine Microsecond Molecular Dynamics Simulations with AMBER on GPUs. 1. Generalized Born. J. Chem. Theory Comput. 8, 1542–1555 (2012).

70. Salomon-Ferrer, R., Götz, A. W., Poole, D., Le Grand, S. & Walker, R. C. Routine Microsecond Molecular Dynamics Simulations with AMBER on GPUs. 2. Explicit Solvent Particle Mesh Ewald. J. Chem. Theory Comput. 9, 3878–3888 (2013).

71. Lindorff-Larsen, K. et al. Improved side-chain torsion potentials for the Amber ff99SB protein force field. Proteins 78, 1950–8 (2010).

72. Jorgensen, W. L., Chandrasekhar, J., Madura, J. D., Impey, R. W. & Klein, M. L. Comparison of simple potential functions for simulating liquid water. J. Chem. Phys. 79, 926 (1983).

73. Meagher, K. L., Redman, L. T. & Carlson, H. A. Development of polyphosphate parameters for use with the AMBER force field. J. Comput. Chem. 24, 1016–25 (2003).

74. Darden, T., York, D. & Pedersen, L. Particle mesh Ewald: An N.log(N) method for Ewald sums in large systems. J. Chem. Phys. 98, 10089 (1993).

75. Eastman, P. et al. OpenMM 4: A Reusable, Extensible, Hardware Independent Library for High Performance Molecular Simulation. J. Chem. Theory Comput. 9, 461–469 (2013).

76. Noé, F. & Clementi, C. Kinetic distance and kinetic maps from molecular dynamics simulation. J. Chem. Theory Comput. 11, 5002–5011 (2015).

77. Mcgibbon, R. T. et al. MDTraj: a modern, open library for the analysis of molecular dynamics trajectories MDTraj: a modern, open library for the analysis of molecular dynamics trajectories. Biorxiv.Org 0-2 (2014). doi:10.1101/008896

78. Harrigan, M. P. et al. MSMBuilder: Statistical Models for Biomolecular Dynamics. Biophys. J. 112, 10–15 (2016).

79. Pérez, F. & Granger, B. E. IPython: a System for Interactive Scientific Computing. Comput. Sci. Eng. 9, 21–29 (2007).

80. Hunter, J. D. Matplotlib: A 2D graphic environment. Comput. Sci. Eng. 9, 90–95 (2007).

81. Pedregosa, F. & Varoquaux, G. Scikit-learn: Machine learning in Python. J. Mach. Learn. 12, 2825–2830 (2011).

82. Amitabh Varshney, Frederick P. Brooks, Jr., Jr. William, W. V. W. Linearly Scalable Computation of Smooth Molecular Surfaces. IEEE Comput. Graph. Appl. 14, (1994).

83. Pérez-Hernández, G. et al. Identification of slow molecular order parameters for Markov model construction. J. Chem. Phys. 139, 15102 (2013).

84. Metzner, P., Schütte, C. & Vanden-Eijnden, E. Transition path theory for markov jump processes. Multiscale Model. Simul. 7, 1192–1219 (2009).

85. E, W. & Vanden-Eijnden, E. Transition-path theory and path-finding algorithms for the study of rare events. Annu. Rev. Phys. Chem. 61, 391–420 (2010).

